# Cell-based model shows complex rearrangement of tissue mechanical properties are needed for roots to grow in hard soil

**DOI:** 10.1101/2022.03.21.485206

**Authors:** Matthias Mimault, Mariya Ptashnyk, Lionel Dupuy

## Abstract

When exposed to increased mechanical resistance from the soil, plant roots display non-linear growth responses that can not be solely explained by mechanical principles. Here, we aim to investigate how changes in tissue mechanical properties are biologically regulated in response to soil strength. A particle-based model was developed to solve root-soil mechanical interactions at the cellular scale, and a detailed numerical study explored factors that affect root responses to soil resistance. Results showed that growth through increasing soil strength is maintained through the softening of cell walls at the tip, a response likely linked to soil cavity expansion. The model also predicts the shortening and decreased anisotropy of the zone of cell elongation, which may improve the mechanical stability of the root against axial forces. The study demonstrates the potential of advanced modeling tools to help identify traits that confer plant resistance to abiotic stress.

## Introduction

Root growth results from two opposing forces (Figure 1 A). On one hand, gradients in osmotic potential drive water into the cytoplasm and cause a build-up of turgor pressure (Kolb, et al., 2017). On the other hand, tension in the cell wall (Cosgrove, 2016), friction forces at root surfaces (Mackenzie, et al., 2013) and compression from the soil (Colombi, et al., 2017) oppose the turgor pressure and determine cell elongation. Turgor pressure is also affected by factors such as soil matric potential and the hydraulic conductivity of the tissue (Bengough, et al., 2011). The biophysical mechanisms of root-soil interaction are now well formalized. The build-up of turgor pressure in cells results from the mechanics of water and ion transport through the cell membrane (Landl, et al., 2017), whilst the cell wall deformation is defined classically using viscoplastic models (Lockhart, 1965). The soil water content is well described by soil mechanical and hydraulic models such as Mohr Coulomb and Richard’ s equations (Dupuy, et al., 2007).

**Figure 1:**
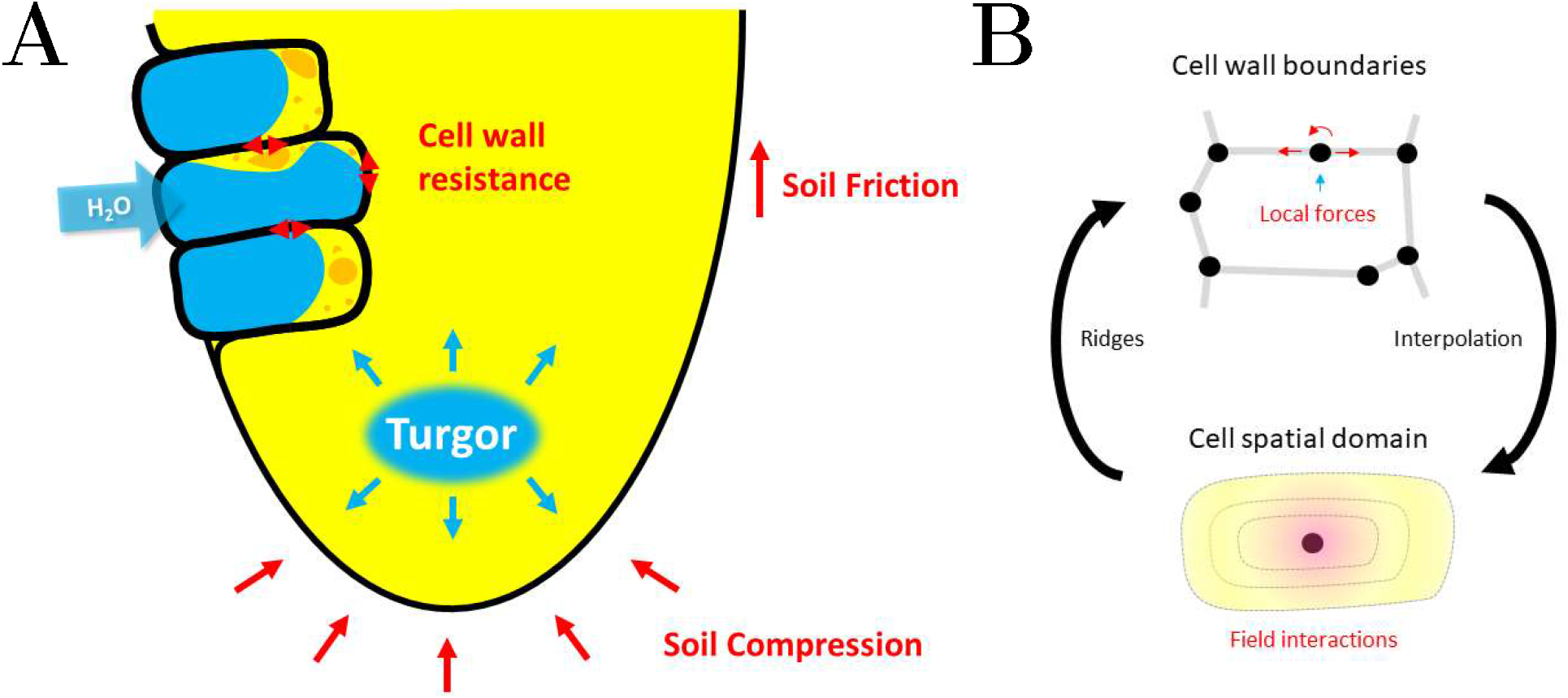
Biophysics of root elongation in soil. (A) Root elongation occurs when turgor pressure exceeds the mechanical resistance of cell walls and the friction and compression from the soil. (B) In cellular models of plant tissues, cells are divided into segments and the balance of forces is computed on each vertex (top). A simplifying approach is to represent the cell using a particle model where cell-cell interactions are computed through a kernel function that weights the effect of individual cells (bottom).

Yet, root responses to soil mechanical resistance are not well predicted by models. Experiments consistently record maximum pressures exerted by a growing root at approximately 1 MPa (Clark, et al., 2003), but roots are observed growing in soils whose mechanical resistance is measured well over 5 MPa (Bengough, 2012). The models developed from first principles by (Dexter, 1987) concluded that an osmoregulation against the soil resistance or the matric potential were needed to predict adequate root elongation rate but, since then, experiments by (Clark, et al., 2001) showed turgor pressure increased by approximately 0.2 MPa when roots are arrested or strongly impeded. It has been suggested that gaps between turgor and soil pressures are due to an overestimation of the friction coefficient of root-soil interfaces from penetrometer resistance tests (Mackenzie, et al., 2013), but experiments show non-linear growth responses to soil resistance (Croser, et al., 1999; Kirby & Bengough, 2002) that are incompatible with the linear response expected from a friction process.

Current limitations in modeling are due to a lack of understanding on how developmental or morphological responses link to adaptation to mechanical stress from the soil. In hard soils, the size of the root elongation zone is reduced (Croser, et al., 1999). The root diameter increases significantly with soil strength, purposely to reduce axial stress and prevent buckling (Martins, et al., 2020). The shape of root tips has also been associated with the ability to overcome mechanical resistance of soils (Colombi, et al., 2017). But to date, it has not been possible to quantify how most of these traits link to a reduced mechanical resistance from the soil. Soils are granular media and particles exert forces that are discrete and heterogeneous. They create opportunities for plant roots to exploit paths of least resistance (Kolb, et al., 2017) and it is difficult to quantify how roots can exploit them to overcome macroscopic pressure from the soil.

A main challenge is to develop models able to grasp the complex relations between root traits and soil properties. A suitable model must describe interactions at the microscopic scale (cells, soil particles) but resolve emerging processes at the macroscopic scale, across entire roots and under changing soil conditions (Dupuy, et al., 2018). In this study we have developed a simple and more efficient computational framework based on the Smooth Particle Hydrodynamics method (SPH) to model the root response to mechanical resistance of the soil. The model is able to compute entire root meristems at cell scale identifying numerical particles with biological cells. The study explores via published data the relationships between cell mechanical properties and soil strength and identifies the modification of tissue properties required to explain observed responses.

## Mathematical model

### Tissue level dynamics

Tissue dynamics are modeled using the framework of continuum mechanics for large deformation. This framework describes the conservation of mass and momentum such that

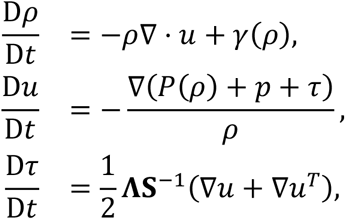

Where 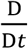 denotes the material derivative. The model equations link the particle velocity *u* (mm h^−1^) to the tissue density *ρ* (μg mm^−3^) and to the Cauchy stress tensor *σ* (MPa), which is the sum of the isostatic stress *P* and a deviatoric component *τ* (shear stress). Here *p* is the difference between turgor pressure and soil pressure and is termed the pressure differential. The matrix **Λ** selects the deviatoric contribution of the compliance tensor required to update *π* such as 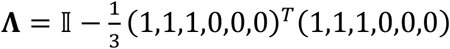.

Growth is modeled through a source term *γ* in the mass conservation equation, given by

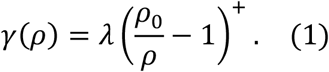

Here *γ* (μg mm^−3^ h^−1^) is expressed as a function of the difference between the equilibrium density *ρ*_0_ and the actual density *ρ*. Hence, the model incorporates a softening mechanism where the material deposition of cell wall is instantaneous. The coefficient *λ* (μg mm^−3^ h^−1^) controls the softening of the cell wall. Only the positive part is used in equation (1) to represent the irreversibility of the extension of the cells. The constitutive equation relating tissue deformation to mechanical stress is given in a weakly compressible setting (Liu & Liu, 2010),

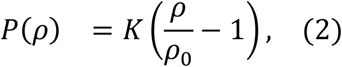

with *K* (MPa) being the bulk modulus. The equations (1) and (2) describe irreversible growth with a densification-relaxation process, similar to the Lockhart and Ortega models (Lockhart, 1965; Ortega, 1985).

To account for the anisotropic deformation of root tissues, the material is assumed to be transversely isotropic in the axial direction, and the compliance tensor **S** is defined using the Voigt notation,

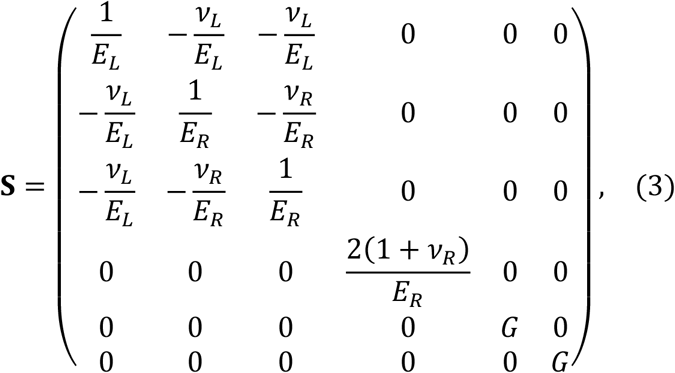

where *E*_*L*_ and *E*_*R*_ are axial and radial moduli, respectively, *v*_*L*_ and *v*_*R*_ are the Poisson moduli in the tangential and radial planes, respectively, and *G* is the shear modulus in the axial direction. Finally, using equation (3), we obtain the expression for the bulk modulus

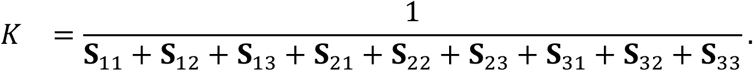

### Cell level dynamics

The individual cells in the tissue are described by particles carrying state variables such as mass, physical dimensions and velocity. The cells are described as ellipsoids (Figure 2 A), by a symmetric positive definite matrix *Q* such that

**Figure 2:**
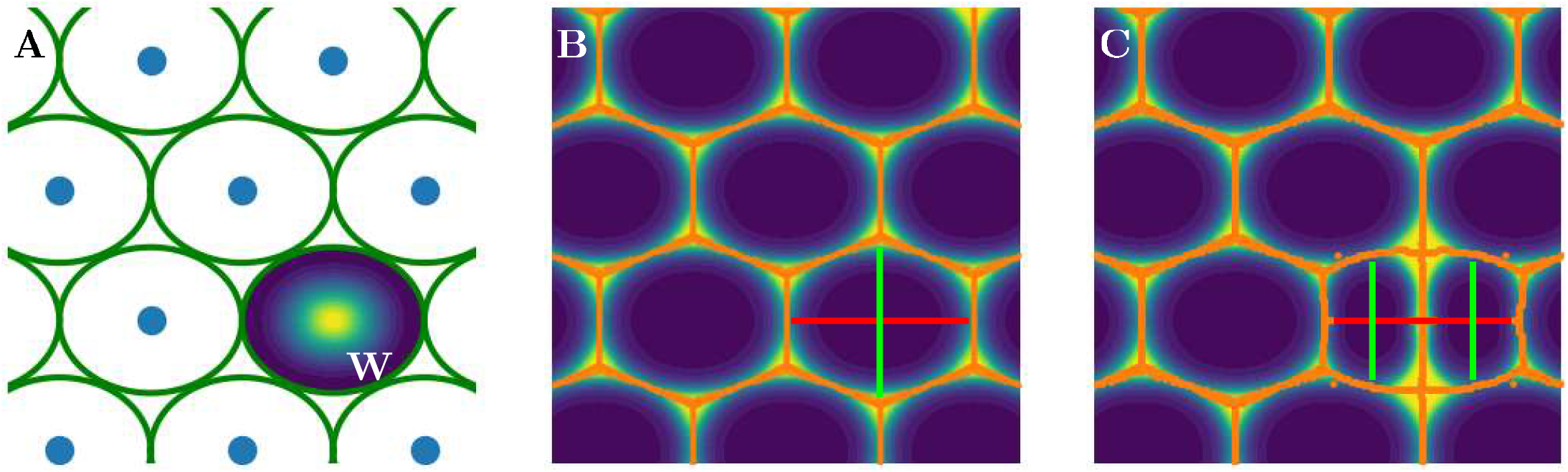
The cell model. (A) Cells within tissues are defined as ellipsoidal particles carrying field variables. The interaction between cells is described through the kernel function *W* associated with the particle (see Supplementary Method), which is a function that decreases with the distance from the particle center. (B) Ellipsoids encode the principal axes of the cell but also generate a potential map that defines the location of cell walls (background color, yellow is the probability location and orange is the ridge interpolation of cell walls). (C) When a cell reaches a length threshold, the division occurs along the principal axis of the cell (red), with the creation of two smaller cells.

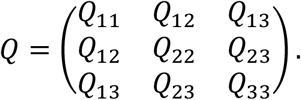

The length of the cell’ s main axes is obtained through the computation of *Q*’ s eigenvalues and eigenvectors,

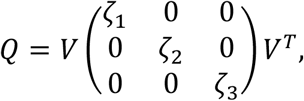

where *ζ*_*i*_ are the eigenvalues and *V* is the matrix composed of the corresponding eigenvectors (Figure 2 B). By convention, *ζ*_*i*_ and *V* are organized such that *ζ*_1_ > 0 is the smallest eigenvalue.

For deformation and division, the state of the ellipsoid is evaluated at discrete time steps noted *t*^*n*^. The deformation of the ellipsoid is updated at every time step *t*^*n*+1^, based on the velocity gradient ∇*u* so that the deformed ellipsoid is defined by *Q*^*n*+1^ for *Δt* = *t*^*n*+1^ − *t*^*n*^

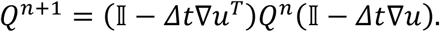

A cell division occurs when the size of the ellipsoid reaches a threshold. Here the cell division threshold 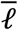 is expressed as a function of the length of the principal cell axis 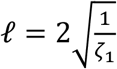 so that division occurs when 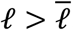. Cell division generates two equal particles on each side of the division plane along the longest axis of the mother particle, at half radius distance (Figure 2 C). Following division at *t* = *t*^*n*^, the updated ellipsoid *Q*^*n*+1^ is then computed as

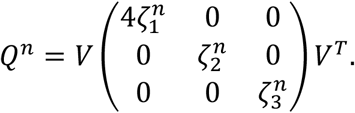

Since the division plane occurs perpendicular to the longest axis, it is consistent with energy minimization principles used in cell division models (Besson & Dumais, 2011).

### Smoothed Particle Hydrodynamics implementation

Unlike classical Finite Element or Finite Volume Methods, mesh-free methods use particles without a predefined topology of neighbors. The SPH approximation of a variable, denoted by ⟨·⟩, is calculated from the weighted interaction of neighboring particles. The weighting is obtained with the kernel function (Figure 2). The kernel function is related to the cell size through the smoothing length coefficient 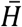, see Supplementary Method. The time evolution of state variables is computed with a prediction-correction algorithm (Crespo, et al., 2015). It computes first a prediction of the evolution at the half time step, and then corrects the estimation at the complete time step. The state variables of particle *i* at half time step 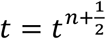 are computed as

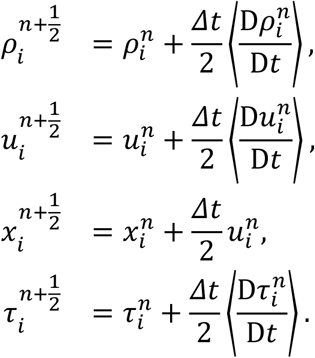

Then at the complete time step *t* = *t*^*n*+1^, state variables are computed as

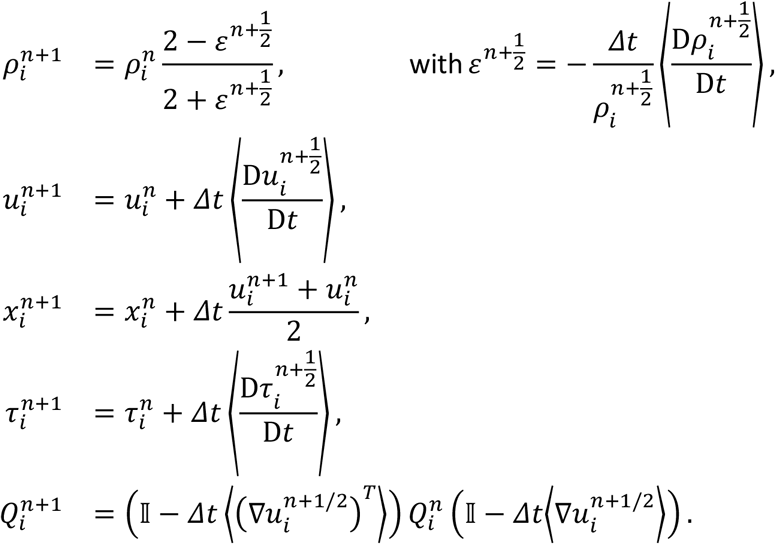

Details on SPH approximation for 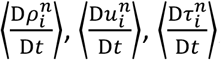, and 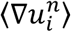 are specified in Supplementary Method.

Computations were performed using initial particle distributions either extracted from microscopy data or generated on a Cartesian lattice. The initial geometries consisted of a rectangle (2D) or cylinder (3D) merged with a half disk or half sphere, respectively. In the absence of specific hypotheses on cell size distribution, particles were initiated as spheres with initial diameter 𝓁 = 𝓁_0_. At the base of the domain, particles were fixed and prevented from moving to represent the connection with the plant body. Particle velocity and mechanical stress was assumed to be zero at initial conditions. Duration of computational time was determined so that the model has reached a steady state. The Courant-Friedrichs-Lewy condition controlling the stability with respect to time step *Δt* is set to 0.9.

The code is implemented in C++ (compiled with v140 tool set on Windows and gcc 8.4 in Linux) and source files are available at https://github.com/MatthiasMimault.

### Parameterization of the model and study cases

Model parameters such as axial/radial modulus, softening coefficient and cell division threshold were assumed to depend on the soil resistance and the axial position *x* along the root, such that

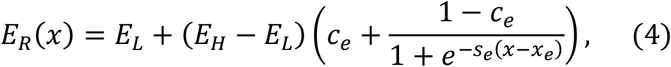

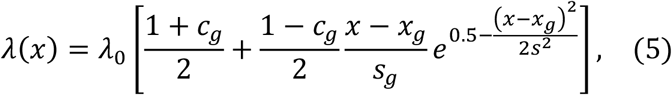

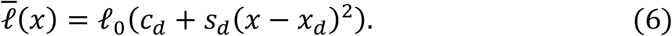

Equation (4) defines the distribution of the radial modulus *E*_*R*_ along the root and causes the deformation to be isotropic at the tip and anisotropic at the base of the root with *E*_*H*_ being a maximal radial modulus. Equation (5) describes the distribution of the softening coefficient *λ* along the root and affects the size and kinematics of the growth zone. Equation (6) describes the distribution of the cell division threshold 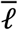 along the axial direction of the root and defines the kinematics of the cell division zone. Model parameters used in numerical simulations are specified in Table 1.

**Table 1:**
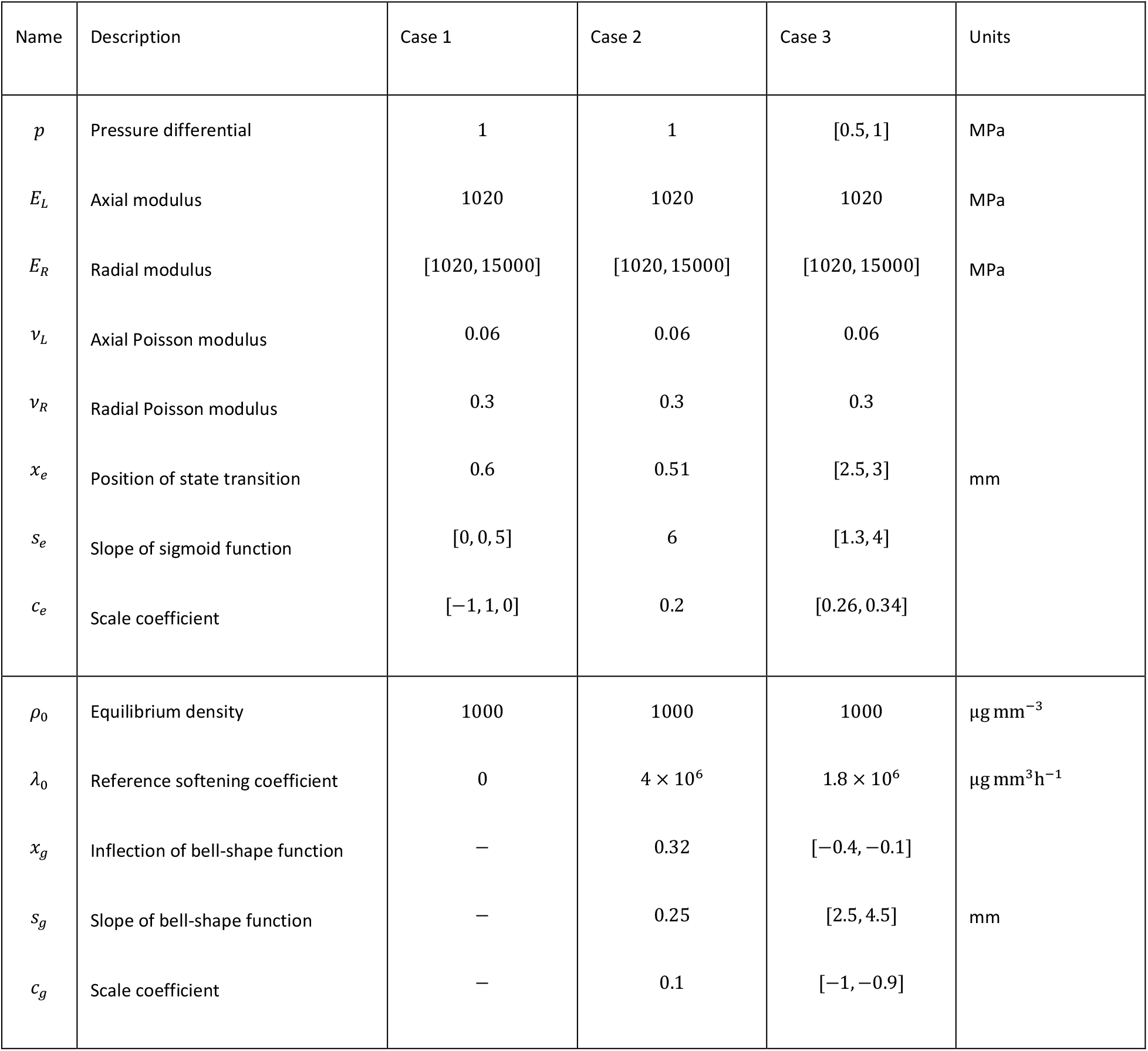
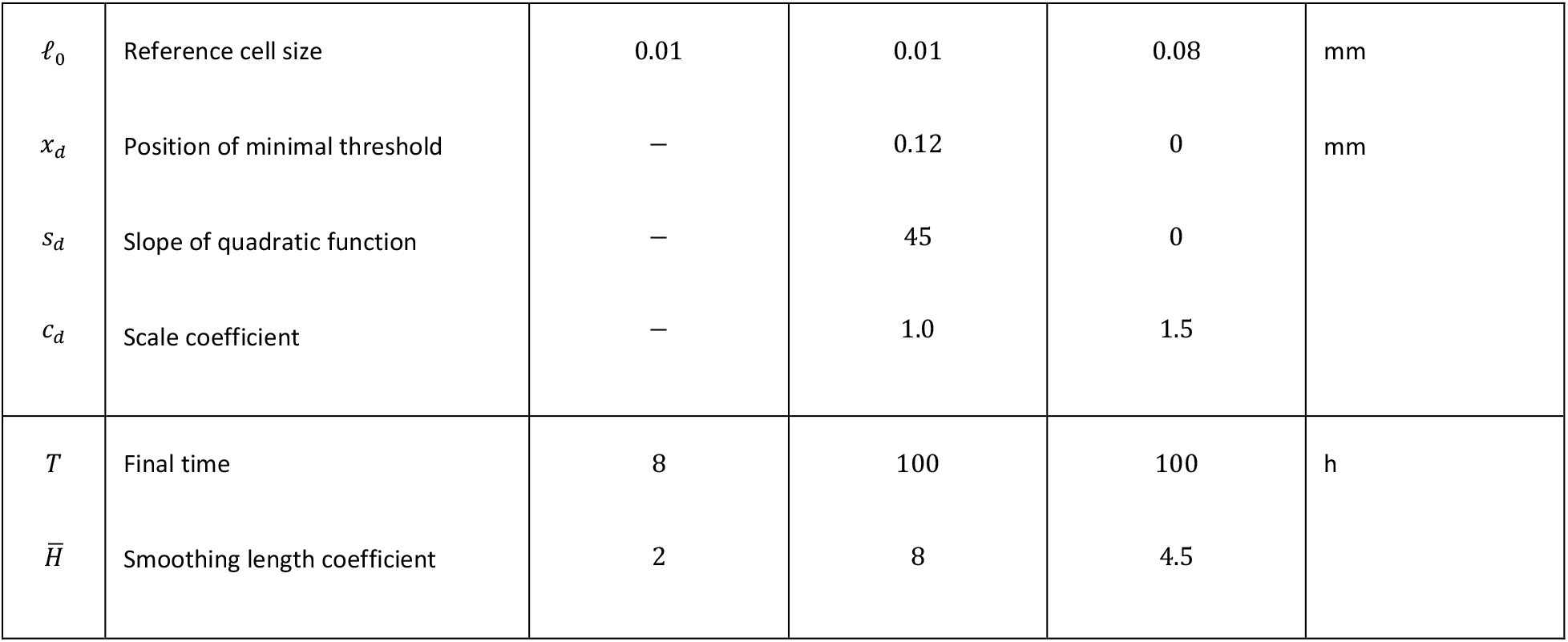
Description of model parameters. When cases involve more than one set of parameters, differing parameters are listed into brackets.

The parameter vector *ξ* = (*p*_*e*_, *c*_*e*_, *s*_*e*_, *x*_*e*_, *p*_*g*_, *c*_*g*_, *s*_*g*_, *x*_*g*_, *c*_*d*_, *s*_*d*_, *x*_*d*_) describes how tissue mechanical and biological properties vary along the root (Table 1). To determine how the root growth adapts to soil condition, first, the parameter vector *ξ*_1_ is determined to predict growth in soft soil conditions *p* = *p*_1_, and then, the parameter vector *ξ*_2_ is determined to predict growth in hard soil conditions *p* = *p*_2_. Finally, the dependency of the root parameters on the soil pressure was defined by interpolating between those two parameter sets following

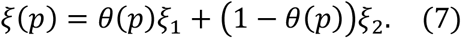

Here, *θ* is a linear function such that *θ*(*p*_1_) = 1 and *θ*(*p*_2_) = 0. Following (Bengough & Mullins, 1991), the calculation of the pressure differential assumed soil resistance to be four times smaller than the penetrometer resistance.

Three study cases are considered to investigate the impact of soil resistance on root growth.

#### Case 1 - Tissue anisotropy and the morphology of the root tip

A remarkable property of developing roots is their ability to maintain their shape and size during growth. To investigate the ability of our model to exhibit such behavior, we performed simulations considering three possible expressions for the radial modulus *E*_*R*_ in equation (4). Constant isotropic properties were modeled using *c*_*e*_ = −1 and *s*_*e*_ = 0. Constant anisotropic properties were modeled using *c*_*e*_ = 1 and *s*_*e*_ = 0. Regulated properties were modeled using parameters shown in Table 1. To assess the ability of the model to correctly predict anisotropy, we considered the case of constant stress in an anisotropic material so that the elongation *δ* of the tissue could be determined analytically.

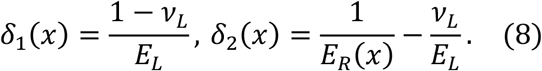

The solutions of the model can then be compared to numerical solutions with elongation estimated with the difference between positions at equilibrium times and initial times.

#### Case 2 - Growth in unimpeded conditions

In the second study case, we examined the ability of the model to predict the unimpeded growth of the root. The model was adjusted to the data proposed by (Beemster & Baskin, 1998) for the spatial profile of cell length, cell elongation and division rates. For the model to accurately predict the growth kinematics parameters, we adjusted the radial modulus, the softening parameter, and the division threshold in equations (4-6). We considered the first 0.6 mm of the root tip because it is where most cell division occurs.

#### Case 3 - Growth in impeded conditions

In the final study case, we examine how the root biophysical parameters vary as a function of soil strength. The model was adjusted to the data proposed by (Croser, et al., 1999) for elongation rate and (Kirby & Bengough, 2002) for root radius to predict the measured elongation rates and root radius for plants grown in sand and clay loams for different soil penetrometer resistance. The pressure differential is determined as the difference between a turgor pressure of 1 MPa and one fourth of the measured penetrometer resistance, so that *p* varied from 1 (soft soil) to 0.5 (hard soil).

### Data processing and Segmentation

To compute statistics from SPH simulations, particle data were aggregated into bins of equal lengths. For a given time step, particles located in the same bin are used to compute the mean and the standard error (SE). The division rate is computed as the ratio between new particles and old particles during the time step. Mean and standard error are computed for each bin at 40 different time steps. The fit between simulations and experimental data was evaluated with the Relative error (*RE*), the Pearson coefficient of determination (*R*^2^) and the Rounded mean square error (*RMSE*), computed with the *sklearn* package (version 0.24.1) in python (version 3.8.8).

To perform computations on realistic cellular architecture we used image stacks with MorphographX (de Reuille, et al., 2015). Results of the image analysis (.csv files) were used as input for SPH computations. The files contained a unique cell identifier, spatial position and ellipsoid approximation for each cell. Results of SPH computations are exported as .vtk and .csv files to be visualized using Paraview (Ahrens, et al., 2005). The cell walls were rendered using a local minima search algorithm (find_peaks, scipy version 1.6.2) applied to the kernel function potential (Figure 2).

## Results

### Coordination of radial and longitudinal growth is needed to maintain the shape of the root apical meristem

Results showed that the distribution of radial modulus *E*_*R*_ along the root has a strong effect on the morphology of the tip (Figure 3). On one hand, when *E*_*R*_ was equal to *E*_*L*_, isotropic expansion was observed at the tip, which results in a spherical outgrowth (Figure 3 A, left). On the other hand, when *E*_*R*_ was significantly larger than *E*_*L*_, uni-axial expansion was observed, transforming the parabolic shape of the root tip into an increasingly sharp structure (Figure 3 A, right).

**Figure 3:**
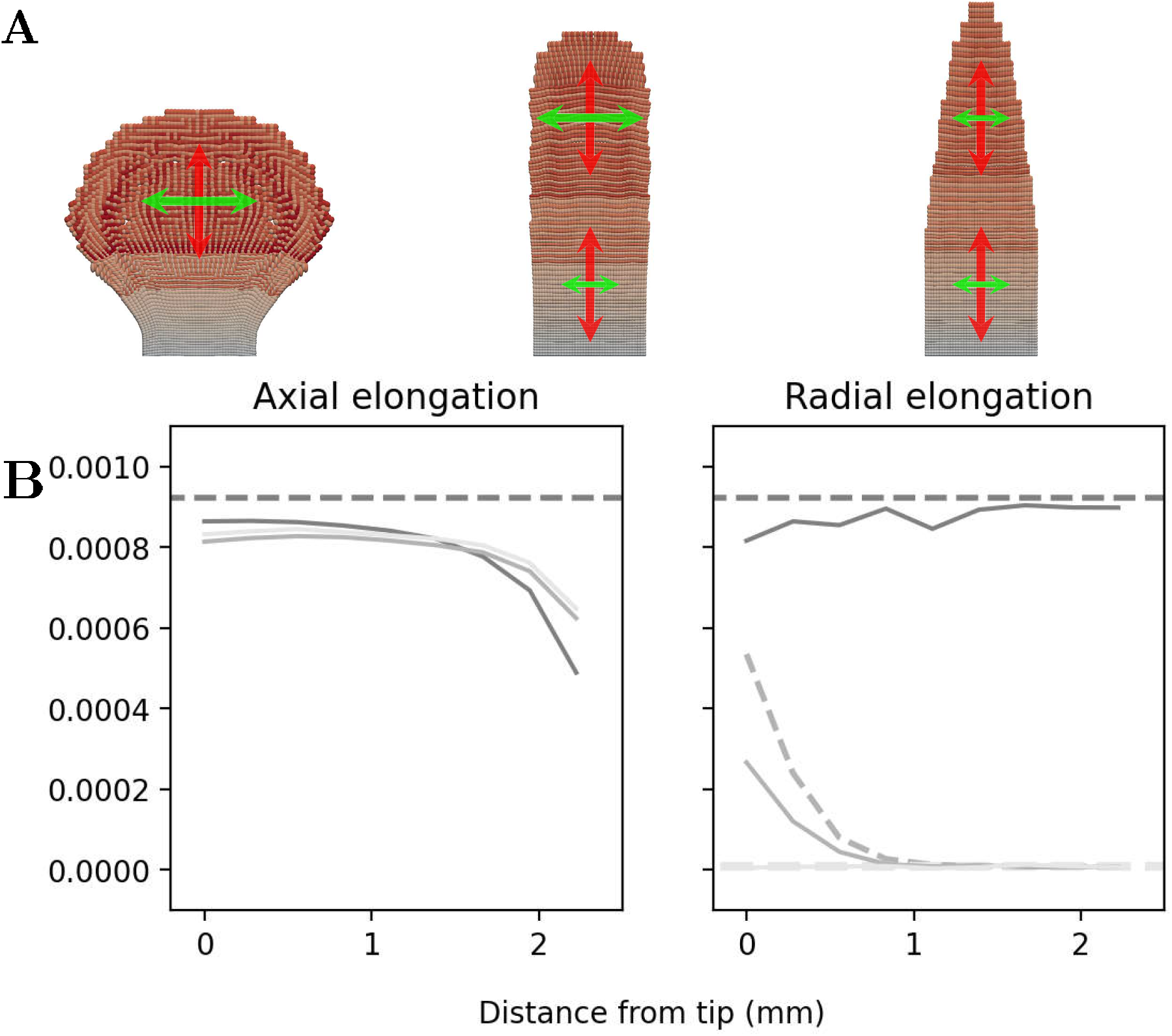
Impact of anisotropy of mechanical properties on the formation of root tip. (A) Three cases were used to test the ability of the model to describe root morphogenesis. The isotropic case (left) was modeled using a radial modulus *E*_*R*_ equal to the longitudinal modulus *E*_*L*_. The anisotropic case (right) was modeled using *E*_*R*_equal to the maximal radial modulus *E*_*H*_. The regulated case (center) was modeled using a radial modulus *E*_*R*_ increasing smoothly from *E*_*L*_ at the tip to maximum radial modulus at the base, using parameters shown in Table 1. Arrows describe the resulting axial elongation *δ*_1_ (red) and radial elongation *δ*_2_ (green). (B) Comparison of SPH predictions with analytical solutions for the isotropic, anisotropic and regulated cases for axial elongation (left) and radial elongation (right). Dashed curves represent analytical solutions and solid curve numerical results. The isotropic, regulated and anisotropic cases are represented in dark to light grey, respectively.

We found that the distribution of the radial modulus *E*_*R*_ that best preserves the shape of the root tip was a sigmoid function in equation (4), specifying the transition from the cell division zone to the cell elongation zone (Figure 3 A, center). The inflection point *x*_*e*_ was found at 0.6 mm so that the transition occurred within half a mm from the root tip. The slope *s*_*e*_ was found to be 5 and *c*_*e*_ was 0.

The accuracy of the model was tested for the isotropic, anisotropic and regulated cases (Figure 3 B). Axial deformation was expected to be constant along the root in all three cases, and the SPH model predicted well the level of axial deformation close to the tip (RE ~ 10%, *x* < 1.5mm). The accuracy of deformation prediction reduced slightly at *x* = 2 mm, which is the location of boundary particles. Since particles are fixed at the boundary, deformation of their neighbors was constrained. The SPH model predicted well the radial deformation for the isotropic case (RE ~ 5%). The accuracy of the prediction in the regulated case was reduced due to the smaller number of particles at the tip (RE ~ 33%). The accuracy in the anisotropic case was lower because deformations were very small and the model was more susceptible to numerical errors (RE ~ 24%).

### Root development in unimpeded conditions

We identified parameters that best fitted experimental data for cell elongation rate, cell length and cell division rate. Results showed the softening of cell walls must increase from the root tip following a bell function with a maximum located at *x*_*g*_ = 0.15 mm in equation (5). We found that the cell length threshold 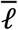 for cell division followed a quadratic function with a slope *s*_*d*_ = 26, a minimum located at *x*_*d*_ = 0.15 mm, and scaling coefficient *c*_*d*_ = 1.5 in equation (6). To maintain stable root diameter and tip shape, we found that the radial modulus *E*_*R*_ must change sharply at the inflection point *x*_*e*_ = 0.15 mm. The size of cells varies greatly on the first 0.6 mm of the root tip. The model uses a smoothing length coefficient 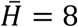, which provides sufficient stability and computational efficiency. Larger cells were more difficult to include in a computationally efficient way because larger 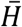 values required for numerical stability would over-smooth dynamics at smaller scales.

Using these parameters, stability is obtained for simulations of long durations. For example it was possible to predict the growth of roots for 10 hours, during which the root volume increased by a factor of 10 (Figure 4 A). 2D computations were performed in 12 minutes using a computer with 32 cores. Predicted elongation rate matched well the values measured experimentally by (Beemster & Baskin, 1998). The elongation rate and cell length distribution were predicted best (Figure 4 B & C) with R^2^ coefficient above 99 % for both and RMSE being equal to 6.2 × 10^−7^ and 0.35, respectively. The cell division rate was better predicted by the model in the root body than at the tip (Figure 4 D). The division rate estimation is poorer whilst the error remains low in the elongation region (R^2^ ~ 18 %, RMSE ~ 1.3 × 10^−3^), From the tip to 0.2 mm, the simulated division rate was much higher than in experimental observations due to radial expansion whilst showing good agreement with observation from 0.2 mm to 0.5 mm.

**Figure 4:**
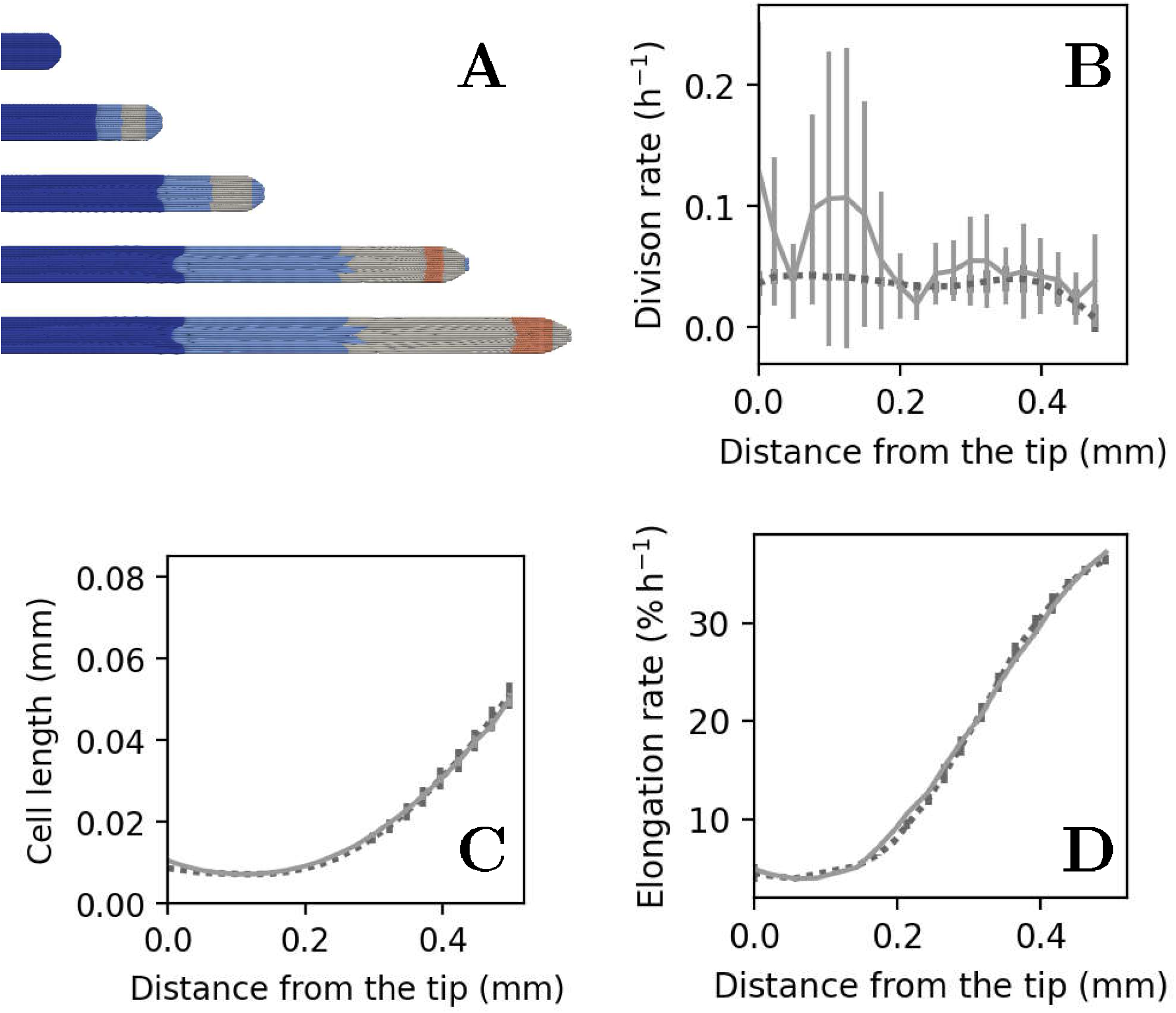
Growth in unimpeded conditions. (A) Simulation of the growth of a root for 10 hours. Colors indicate the number of cell divisions, blue (0) to red (3) (visible on web version). (B) Comparison between experimental and SPH predictions for cell division rate. (C) Comparison between experimental and SPH predictions for cell lengths. (D) Comparison between experimental and SPH predictions for cell elongation rate. Symbols are mean ± SE. Observations are represented by dark dashed lines and numerical solutions by solid light lines.

### Root response to soil mechanical resistance

The final series of simulations were performed to explain how root mechanical properties change in response to increase of soil strength. We identified parameters in equations (4-6) that best predicted experimental distribution for cell elongation rate and axial distribution of root radius. From the tip to 1 mm, the elongation rate of roots is observed to remain the same across the range of pressure differential. The size of the root elongation zone however varied with soil strength. In loose soil, the size of the elongation zone was larger and the root elongation rate was higher and occurring on a wide region of the tip, with a maximum further away from the root tip. The softening coefficient was found to increase closer to the tip and have a tail that declines faster with the distance from the tip (Figure 5 A). Growth parameters in equation (5) depend on the pressure differential *p* using equation (7) with parameters *x*_*g*_ = −0.4 mm, *s*_*g*_ = 4.5 and *c*_*g*_ = −1 for *p*_*g*_ = 0.987 MPa, and *x*_*g*_ = −0.1 mm, *s*_*g*_ = 2.5 and *c*_*g*_ = 0.9 for *p*_*g*_ = 0.625 MPa, and *c*_*g*_ = 0. Results of simulations showed the increased softening at the root tip was sufficient to predict accurately the changes in cell elongation rate observed experimentally (Figure 5 B).

**Figure 5:**
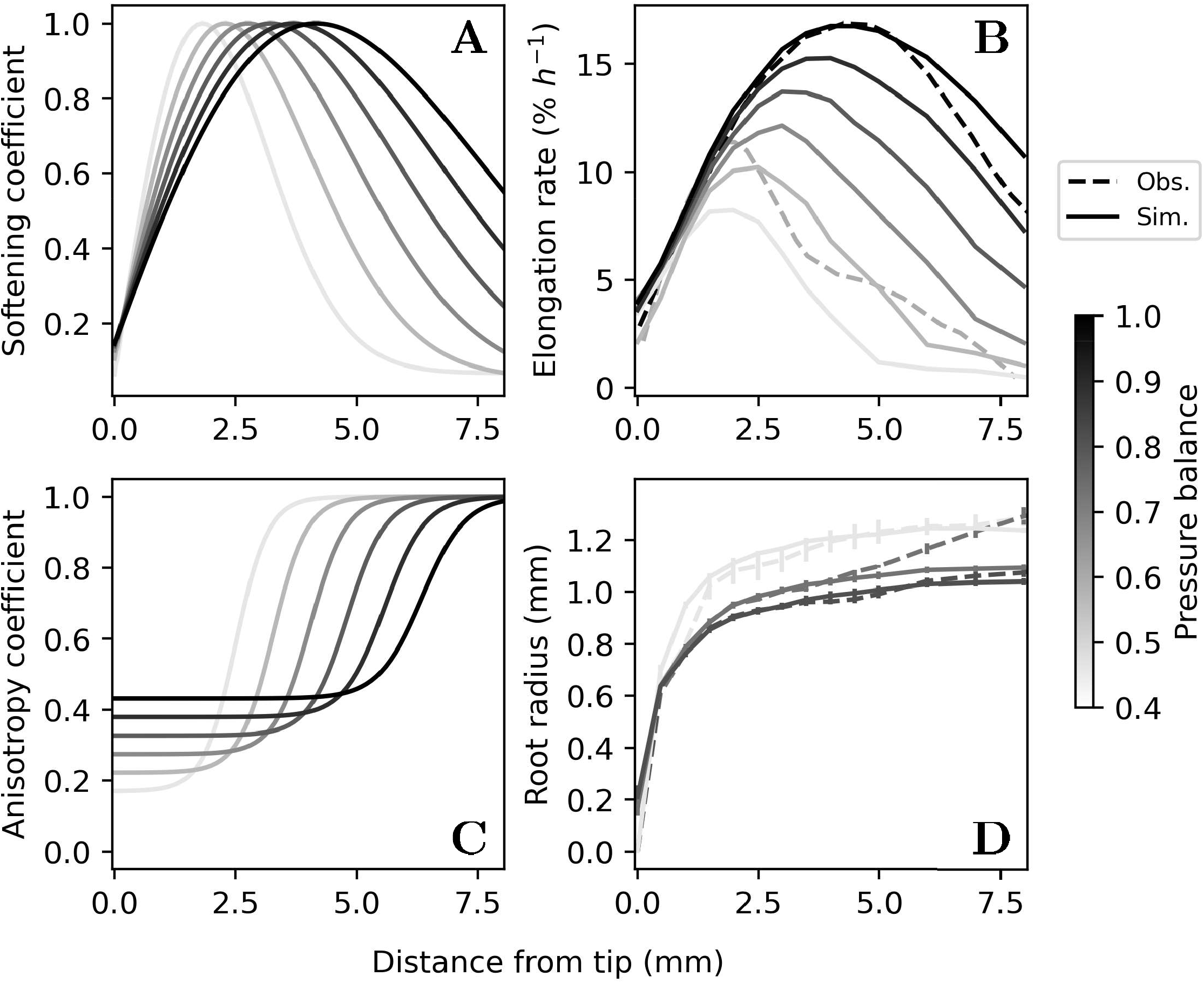
Changes of cell wall softening and anisotropy coefficients in response to increases in soil strength. (A) The softening coefficient λ was found to reach a maximum closer to the tip and tail off quicker. (B) This distribution allowed the SPH model to match the elongation rate observed experimentally. (C) The radial modulus *E*_*R*_ was found to take lower values in a smaller region. (D) This distribution of anisotropy allowed the SPH model to match the profile of root radius observed experimentally. Observations are represented by dashed lines and numerical solutions by solid lines. Increase in the pressure differential is represented from bright grey to dark.

To predict the increase in root radius, the radial modulus *E*_*R*_ must also vary with the pressure differential *p*. We found the parameters that best predict the radial expansion of the roots were *x*_*e*_ = 3 mm, *s*_*e*_ = 1.3 and *c*_*e*_ = 0.34 for *p*_*e*_ = 0.825 MPa, and *x*_*e*_ = 2.5 mm, *s*_*e*_ = 4.0 and *c*_*e*_ = 0.26f or *p* = 0.5 MPa. These parameters resulted in a shift of the anisotropic transition closer to the root tip and a smoother transition through the elongation zone. Close to the tip, the root tissue was also more isotropic in hard soil (Figure 5 C).

Using these parameters, the SPH model could predict accurately the increase in root diameter observed experimentally (Figure 5 D). The model predicted well the 20% increase in root diameter in the mature part of the root with a continuous shift following the change in pressure differential (*R*2 > 98 %, *RMSE* < 0.004 for *p* = 0.5 and 0.825 and *R*2 > 97 %, *RMSE* < 0.007 for *p* = 0.75). Elongation rates are well reproduced (*R*2 ~ 86 %, *RMSE* ~ 1.6 for *p* = 0.987 and *R*2 > 97 %, *RMSE* < 0.78 for *p* = 0.625). Discrepancies between experimental observations and predictions occurred in the mature region of the root. These were due to the fact that the model only accounted for development at the root apical meristem and no softening was allowed beyond that point of the root.

## Discussion

### New framework for modeling root meristems at cell scale

There are only a handful of modeling approaches able to resolve biophysical interactions within a growing root at cell scale. Early tissue models have adopt 1D continuous description (Dexter, 1987; Ortega, 1985) and have exposed the balance of forces acting on growing roots. Since then, models have included more detailed endogenous (Chavarría-Krauser & Schurr, 2004) and exogenous processes (Dupuy & Silk, 2016), but the use of one dimensional descriptions remains limiting. For example, it is not easy to describe bending and buckling caused by rigid obstacles, or growth through path of least resistance in soil (Kolb, et al., 2017). The extension to higher dimensions proposed by (Hejnowicz & Karczewski, 1993) have met with limited success because it requires many analytical calculations. But numerical techniques such as the Finite Element Method (FEM) have considerably advanced the understanding of biophysics of root growth. Applications now include predictions of mechanical stress and failure zone in soil, water transport in roots, or studies of root gravitropic responses (Kirby & Bengough, 2002; Grieneisen, et al., 2007; Dupuy & Silk, 2016).

Cellular models have emerged subsequently as a mean to account for morphogenetic processes. The use of vertex approaches has proven particularly powerful to solve complex developmental mechanisms, including phyllotaxis, lateral root formation, cell differentiation patterns on leaves, and cell-cell communication (Smithers, et al., 2019). Recent computational techniques couple vertex dynamics to cell volume and implement quasi static approximations to improve the stability of computations. However, because each cell in the system must be described through a large number of vertices, computations are slow and difficult to parametrize and applications have largely focused on problems with reduced physical dimensions, e.g. outermost layers of cells, or using two-dimensional descriptions (Grieneisen, et al., 2007; Fozard, et al., 2013; Bidhendi & Geitmann, 2017).

Particle based approaches are relatively recent alternatives. Numerical approximations are less well characterized than finite element methods, and the calibration of numerical algorithms remains complex (Vacondio, et al., 2021). The approximations computed by the Smoothed Particle Hydrodynamics method depend strongly on the particle distribution. Furthermore, current software or libraries are community led and usually problem specific. However, since the first publication of the SPH method in 1977 (Gingold & Monaghan, 1977; Lucy, 1977) the field of research has progressed at a considerable pace with for example the introduction of anisotropic kernels to compute large deformations, implicit and incompressible schemes to stabilize long-term simulations, corrected and symmetric formulations to manage numerical errors, and improved neighbor search for better efficiency (Liu & Liu, 2010). Finally, communities are pushing for development of unified SPH framework using open Application programming interfaces (Zhang, et al., 2021).

Using SPH, we have demonstrated the feasibility of computing root-soil interactions at cell scale, across the entire organ. The technique allowed us to associate numerical particles with biological cells, preserving the shape, size and cellular architecture of the root with a minimal number of degrees of freedom. The vertex and particle modeling approaches are therefore dual descriptions of the cellular architecture (Figure 1 B). Particles describe the interior of the cell, and cell walls can be reconstructed from the kernel function (Figure 2). Flexible numerical schemes allowed computation of thousands of cells using a modern workstation (Mimault, et al., 2019). The SPH method proved amenable to high dimensional and multi-physics problems, tests performed on two-dimensional domains could be easily ported to three dimensional problems without the need for a new interpolation scheme (Supplementary Video 3 and 4). Simulations could be performed using images of the root anatomy with limited image processing because only basic shape descriptions for the cells are needed as an input (Figure 6).

**Figure 6:**
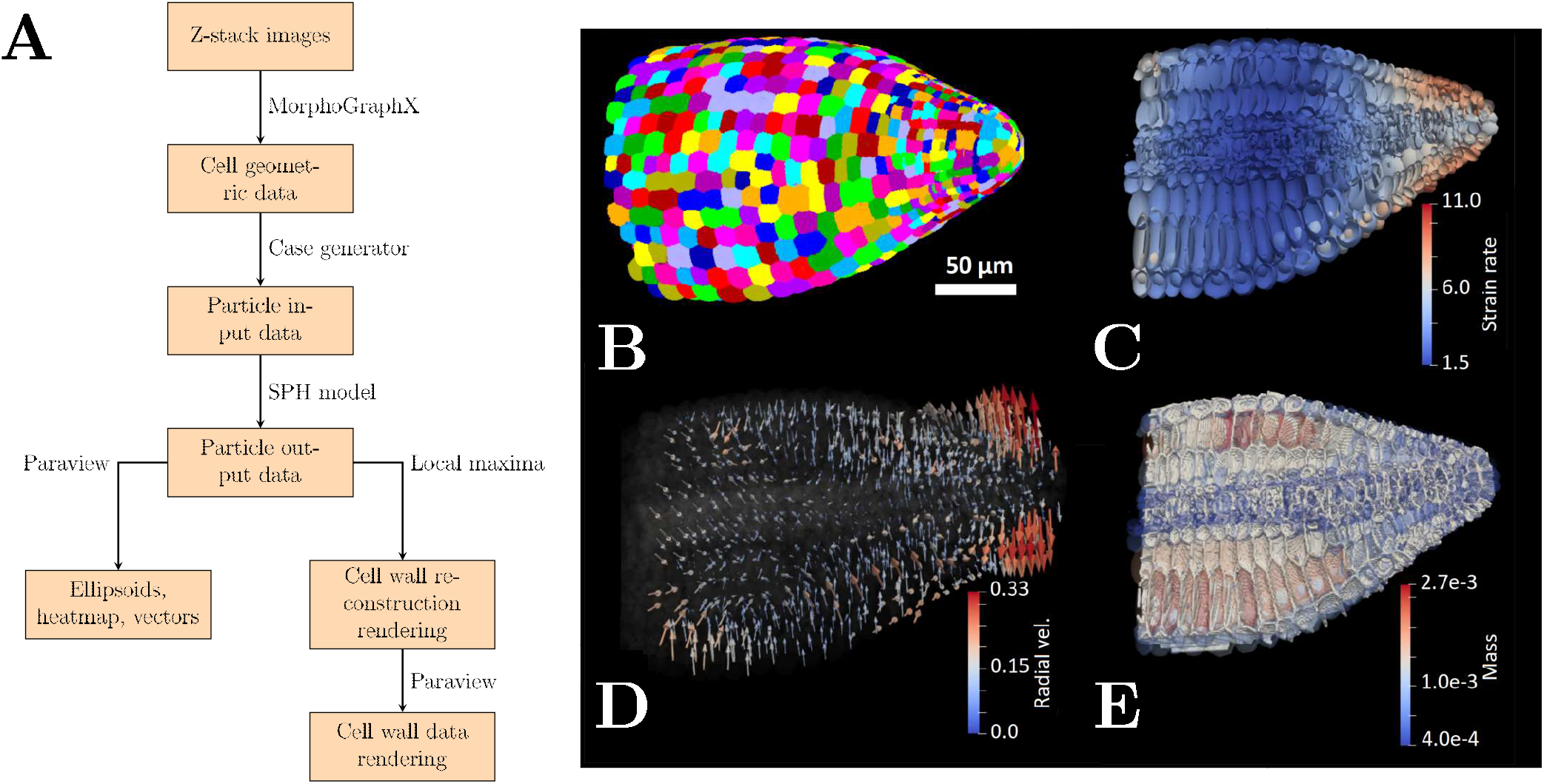
Software pipeline for using microscopy data as input for SPH simulations. (A) The software pipeline is developed to carry out SPH computations including pre and post processing. (B) The pipeline is used to extract cell geometrical features from 3D microscopy data that is imported as particles carrying physical, biological and mechanical information. The model then simulates the growth of plant roots, including mass accumulation and cell division. Data obtained from simulations include the elongation rate (C); radial cell velocity (D). Kernel functions are also used to visualize cell walls (E).

### Root strategies to overcome mechanical resistance from the soil

Using our SPH model, we could accurately predict the growth of a root from its biophysical parameters (Figure 4). The model assumed a constant turgor pressure because only limited increases in turgor pressure are measured in arrested roots (Clark, et al., 2001). To explain growth rates observed in soils which penetrometer resistance exceed turgor pressure, we choose to represent the pressure perceived by the plant, which can be approximated as a fraction of the penetrometer resistance (Bengough & Mullins, 1991).

The model revealed how tissue mechanical properties are rearranged when soil strength is increased. We first observed that pressure from the soil alone was not able to predict the morphology of roots growing in hard soil. Plant roots are known to reduce the size of their meristem but also to maintain the cell elongation rate at the root apex. In soft soil, the elongation region is large and the root radius rather thin whilst in hard soil, it is the reverse situation, with the elongation concentrated at the tip and a larger radius. The observation was made in response to mechanical stress but is also observed during water stress (Bengough & Mullins, 1991). To predict such responses, cell wall softening must reach a maximum closer to the root tip, accounting for the soil containment with relaxed cell wall resistance. This occurs in a small region because cell wall softening was shown to weaken roots that were exposed to axial forces (Kolb, et al., 2017). The softening of the tip root together with the shortening of the growth and softening zone may therefore reduce the susceptibility of the root to buckling (Clark, et al., 2003). Realistic growth responses were obtained without the need for the softening coefficient to increase. This result indicates excessive softening may be detrimental to the mechanical stability of the root tissue.

Root diameter was also observed to increase by 20% to 60% when penetrometer resistance is doubled (Kirby & Bengough, 2002). Axial forces alone can explain a radial expansion of the root due to incompressibility. For example, an axial stress of 1 MPa with a Young’ s modulus of 100 MPa and a Poisson coefficient of 0.3 explains a radial expansion of 0.3%. To match observations, the model also requires a change in the radial modulus. The tissue must become more mechanically isotropic at the root tip and more anisotropic in the cell elongation zone. This response is also consistent with mechanical stability requirements under increased external pressures. Increased isotropy and cell wall softening at the root tip generates soil cavity expansion, a process which has been shown to reduce the axial load on the root tip (Mackenzie, et al., 2013), which results in the larger diameter measured experimentally. In the elongation zone however, the strengthening of the tissue in the radial direction may reinforce the root against axial stress, and together with the increase in diameter may prevent the root from buckling.

### Models for the next generation of root microscopy data

Data available to quantitatively characterize root processes is growing rapidly. Live microscopy in soil-like conditions has greatly improved with the development of artificial soils (Downie, et al., 2012; Ge, et al., 2021). The emergence of light sheet microscopy democratizes the use of instruments built in-house for larger living samples (Liu, et al., 2021). Available techniques to quantify growth, morphology and anatomy from roots grown in natural soils has also improved drastically with techniques such as X-ray tomography, MRI and neutron tomography, and Laser Ablation Tomography (Downie, et al., 2015). Microfluidics and robotics (Massalha, et al., 2017) now enable fast three dimensional reconstruction of the root architectures at anatomical and whole plant level. Likewise, algorithms for processing image data are improving. Software enables larger samples to be reconstructed from a myriad of views (Bria & Iannello, 2012) and the automated extraction and classification of features (Kan, 2017). The availability of software resources to analyze and process root microscopy data is also greatly expanding thanks to repositories for open-source software (Lobet, 2017) and microscopy data (Besson, et al., 2019). The emergence of standards for data formats stimulates portability and communication between software with reduced efforts needed in software development (Pradal, et al., 2008; de Reuille, et al., 2015).

Biophysical models are becoming a critical component of these software pipelines. Models could help forecast root performance in field conditions from quantitative traits now delivered by modern microscopes and live imaging facilities. Currently, mathematical models able to exploit information contained in large imaging data sets are limited, and this paper has demonstrated the potential for particle-based approaches to bridge this gap. The technique requires minimal description of the anatomic structure of the roots, it is computationally efficient and can easily be integrated into pipelines for the simulation of realistic scenarios (Figure 6, and Videos 3 and 4). Results presented here showed that the model and simulation pipeline can reveal the complex responses of roots to the strengthening of soils. The next step now is to confirm the results with a broad range of root anatomy and adequate field validation. This would further consolidate the potential of image based modeling to assist the experimental research.

## Acknowledgment

This project is supported by the consolidator fellowship from the European Research Council ERC SENSOILS-647857. The James Hutton Institute received support from the Scottish Government Rural and Environment Science and Analytical Services Division (RESAS, Workpackage 1.1.1,2.1.6,2.1.7,2.3.4).

## Contributors’ statement

M.M. developed the model, designed experiments and performed computations. L.D. and M.P. supervised the findings. M.M. wrote the manuscript. All authors contributed to the revision of the manuscript.

